# Expression-Driven Genetic Dependency Reveals Targets for Precision Medicine

**DOI:** 10.1101/2024.10.17.618926

**Authors:** Abdulkadir Elmas, Hillary M. Layden, Jacob D. Ellis, Luke N. Bartlett, Xian Zhao, Reika Kawabata-Iwakawa, Hideru Obinata, Scott W. Hiebert, Kuan-lin Huang

## Abstract

Cancer cells are heterogeneous, each harboring distinct molecular aberrations and are dependent on different genes for their survival and proliferation. While successful targeted therapies have been developed based on driver DNA mutations, many patient tumors lack druggable mutations and have limited treatment options. Here, we hypothesize that new precision oncology targets may be identified through “expression-driven dependency”, whereby cancer cells with high expression of a targeted gene are more vulnerable to the knockout of that gene. We introduce a Bayesian approach, BEACON, to identify such targets by jointly analyzing global transcriptomic and proteomic profiles with genetic dependency data of cancer cell lines across 17 tissue lineages. BEACON identifies known druggable genes, e.g., *BCL2, ERBB2, EGFR, ESR1, MYC*, while revealing new targets confirmed by both mRNA- and protein-expression driven dependency. Notably, the identified genes show an overall 3.8-fold enrichment for approved drug targets and enrich for druggable oncology targets by 7 to 10-fold. We experimentally validate that the depletion of *GRHL2*, *TP63*, and *PAX5* effectively reduce tumor cell growth and survival in their dependent cells. Overall, we present the catalog of express-driven dependency targets as a resource for identifying novel therapeutic targets in precision oncology.

## Introduction

Precision oncology requires accurate identification of molecular aberrations in cancer cells that can serve as biomarkers and therapeutic targets. While some tumors harbor genomic mutations predictive of cancer vulnerability, a large fraction of cancer cells lack such actionable mutations^1–3^. Large-scale genetic dependency screens, including the Cancer Cell Line Encyclopedia (CCLE)^4^, Cancer Dependency Map (DepMap)^5^ and CancerGD^6^, have revealed that cancer cells show different vulnerability upon genetic knockdown or knockout. Across diverse types of molecular alterations—including mutations, copy number alterations and expression—gene expression biomarkers have been identified as the top biomarkers of genetic dependency, e.g., in 82% of the 501 DepMap cell lines in a genome-scale RNAi screen^5^. We thus reasoned that precision oncology targets might be identified through “expression-driven dependency”, whereby cancer cells with high expression of the targeted genes are more vulnerable to genetic depletion or therapeutic inhibition.

Multiple studies have used genetic and functional screening data to identify cancer vulnerabilities present in a subset of cancer cells, including aneuploid cancer cells^7,8^, pediatric tumor cells^9^, and multiple myeloma cells^10^. Notable targets identified include the WRN helicase that is essential in cancers with microsatellite instability (MSI) but dispensability in microsatellite stable cells^11,12^, PKMYT1 kinase in CCNE1-amplified tumors, and BCAR1 in KRAS mutant pancreatic cancer, where the suppression of BCAR1 and TUBB3 sensitizes cancer cells to ERK inhibition by reducing MYC protein levels^13^. Bondeson et al.^14^ identified phosphate dysregulation as a therapeutic vulnerability in ovarian cancer through genome-scale CRISPR-Cas9 screens, highlighting the XPR1–KIDINS220 protein complex as crucial for cancer cell survival. Another study^8^ identified the ubiquitin ligase complex UBA6/BIRC6/KCMF1/UBR4 as crucial for the survival of aneuploid epithelial tumors. These studies highlight the potential of developing a systematic approach to identify drug targets by linking subsets of cancer cells to genetic dependency based on their aberrant expression.

Expression analyses focusing on only the transcriptome assume that high gene mRNA expression translates into high protein abundance. However, gene expressions show only moderate correlations with protein expression in cancer cell lines and primary tumors^15–20^, and protein-level analyses may identify new targets^3,21–23^. Notably, global proteomic profiles of 375 cell lines in the CCLE/DepMap were recently generated by global mass spectrometry (MS), quantifying a total of 12,399 proteins using multiplexing quantification methods^24^. The combination of these datasets provides unprecedented opportunities to identify new protein biomarkers and therapeutic targets across cancer types.

Herein, we integrated global proteomic and transcriptomic profiles of 855 cancer cell lines across 17 tissue types from Cancer Dependency Map (DepMap)/Cancer Cell Lines Encyclopedia (CCLE)^24,25^, and the corresponding cancer cell dependency scores (Achilles) based on the CRISPR knockout screens^25–27^. By developing a new Bayesian correlation approach, BEACON, we identified the expression-driven cancer cell dependencies (ED) for each tissue type at different molecular layers, and revealed new potentially actionable targets that are strongly-associated with druggable gene lists^28^ (**Figure 1**). Our analyses identified the known drug targets SOX10 and ESR1 demonstrating strong gene/protein ED linked to their specified cancer type and revealed new potential candidate targets for each cancer type. Experimental validation supported the actionability of the new candidate targets TP63, GRHL2, and PAX5, exposing a vulnerability in their dependent cancer cells.

**Figure 1.**
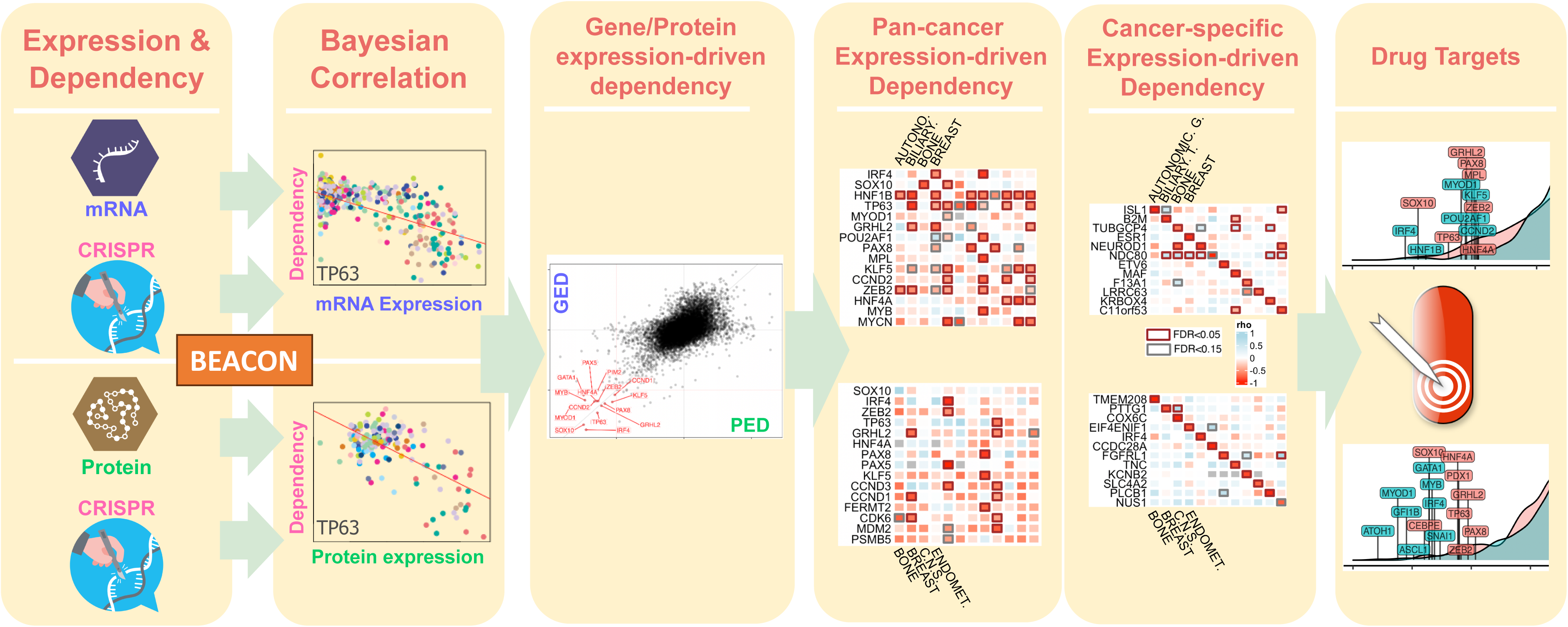
Study overview. (A) The integration of global proteomic and transcriptomic profiles from 375 cancer cell lines across 22 tissue types in the Cancer Cell Lines Encyclopedia (CCLE), with cancer cell dependency scores derived from CRISPR knockout screens (Achilles). (B) BEACON identifies expression-driven dependency (ED) by using a Bayesian estimation of the correlation coefficient between gene/protein expression and cancer cell dependency data across the cell lines for a representative gene (e.g., TP63). (C) Comparison of gene/protein EDs revealed potential markers showing consistency at different molecular levels or arising post-transcriptionally. (D) Heatmaps showing pan-cancer expression-driven dependencies, GED (above) and PED (below), revealing dependencies that are common across multiple cancer types. (E) Heatmaps illustrating cancer-specific expression-driven dependencies, GED (above) and PED (below), identifying dependencies unique to specific cancer types. (F) Identification of new potentially actionable targets that are strongly associated with druggable gene lists catalogued in DrugBank, highlighting their therapeutic potential.

## Results

To identify genes showing expression-driven dependency, we first integrated RNA-seq data, global mass spectrometry proteomics data, and the cell dependency data corresponding to the same cell lines in the DepMap project (**Methods**). We restricted our analyses to lineages where at least 7 cell lines with cancer cell line dependency and corresponding mRNA/protein expression data were available to ensure statistical robustness (**Figure S1A**). Overall, 855 cell lines across the 17 lineages shared cancer cell dependency scores and corresponding mRNA and protein expressions (N=854 for mRNA, N=290 for protein, **Figure S1B, Table S1**). Based on this limited sample size per cell lineage (**Figure S1C)**, we noticed that the basic correlation techniques may lead to spurious correlations, particularly for protein expression (**Figure S1B, Figure S2**). Thus, we developed a Bayesian approach, BEACON (Bayesian EvAluation of expression Correlation-driveN dependency), to model expression levels and dependency scores as the bivariate Gaussians and used Markov Chain Monte Carlo (MCMC) sampling to test the null hypothesis that these two are uncorrelated for each given gene (**Methods**). BEACON offers the unique advantage of utilizing prior distributions that are less susceptible to outliers, especially in multiple lineages where the number of cell lines. We benchmarked BEACON’s Bayesian correlation vs. Pearson correlation by simulating expression and dependency datasets at various correlation levels (from −1 to 1, with 0.25 intervals) and sample size (number of cell lines, 10, 20, 30, 40, 60, 100), with different fraction (0.1, 0.3, 0.5) of samples being outliers (**Figure S2**). Based on these simulations, we observed that the Bayesian method is better than Pearson correlation for estimating moderate true correlation (|rho| < 0.75) in small sample size, and preferable in noisy data (noise level ≥ 0.3, i.e., 30% or more of the samples are corrupted by noise to become outliers), regardless of sample size or true correlation level.

### Cancer vulnerability targets showing gene expression-driven dependency (GED)

We first applied BEACON to reveal cancer vulnerabilities that show gene expression-driven dependencies (GED) at the mRNA level. We first analyzed the pan-lineage GED by using mRNA levels and the corresponding dependency scores from 854 cell lines with available data across 17 lineages and identified 244 genes showing significant association (correlation coefficient, rho < −0.25, FDR < 0.05). The notable genes with strong pan-lineage associations (false discovery rate, FDR < 1e-32) include SOX10 (correlation coefficient, rho = −0.83), IRF4 (rho = −0.82), HNF1B (rho = −0.76), and MYOD1 (rho = −0.70) (**Table S2**).

Having found many strong GEDs across cancer cells from different tissue types, we then applied BEACON to identify tissue-specific GEDs within each lineage (**Methods**). As expected, several significant pan-lineage GED targets also showed substantial tissue-level GED in multiple lineages, including TP63, CCND1, CCND2, and KLF5 (rho ≤ −0.61, FDR < 1e-32) (**Figure 2A**, **Figure 2B**). TP63 showed significant (rho < −0.25, FDR < 0.05) GED across 14 out of 24 lineages of the cancer cell lines. TP63 is a member of the p53-family transcription factors that regulates developmental processes in several organs and tissues, as well as tumorigenesis and tumor progression^29^. Another transcription factor, KLF5 also showed significant (rho < −0.25, FDR < 0.05) GED frequently across half of the cell lineages (12/24). This could be explained by its role in the development and progression of various types of cancer, as its expression is essential for cell cycle regulation, apoptosis, migration, and differentiation, impacting a wide array of target genes such as cyclin D1, cyclin B, PDGFα, and FGF-BP^30^.

**Figure 2.**
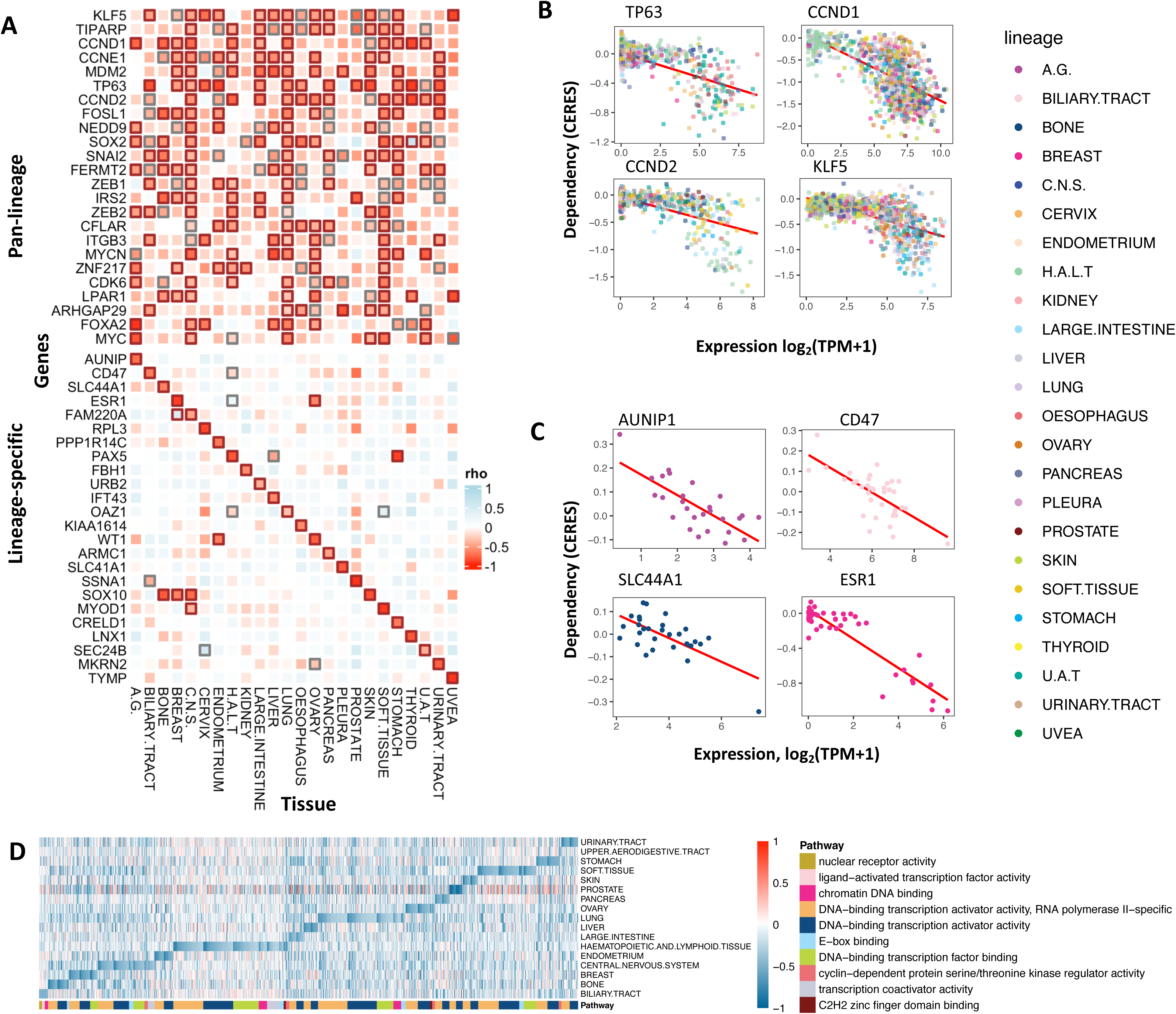
Gene Expression-driven Dependency (GED). (A) Heatmap illustrating pan-lineage and lineage-specific gene expression-driven dependencies (GEDs) across various cancer types. Each square represents the correlation (rho) between gene expression and dependency (CERES scores) in the respective tissue types. Significant dependencies are highlighted with bold outlines (FDR < 0.05 in black, FDR < 0.15 in grey). (B) Scatter plots showing examples of gene expression vs. dependency correlations for selected genes (TP63, CCND1, CCND2, KLF5) with significant pan-lineage dependencies. Data points (cell lines) are colored by tissue type. (C) Scatter plots demonstrating lineage-specific dependencies for selected genes (AUNIP1, CD47, SLC4A1, ESR1). Data points are colored by tissue type, highlighting lineage-specific associations. (D) Pathway enrichment analysis of lineage-specific GEDs, visualized as a heatmap. Each cell indicates the ED score of a particular pathway gene (column) in a specific tissue type (row), with genes grouped (colored) by functional pathways.

Since multiple lineages were dependent on the expression of some transcription factors such as KLF5 and TP63, targeting these genes may lead to unintended consequences across tissue types. To minimize potential off-target effects, we further identified the GED targets showing only lineage-specific expression-driven dependency, i.e., exhibiting low correlation (more negative rho) within a given lineage’s cell lines and relatively smaller (near-zero) correlation in other lineages (**Methods**, **Figure 2A**). Among such targets, we found MYOD1 for soft tissue, PAX5 for haematopoietic and lymphoid tissue, SOX10 for skin, and ESR1 for breast (rho ≤ −0.84, FDR < 1e-32) (**Table S2, Figure 2A**, **Figure 2C**), the latter of which is an already well-targeted gene through hormonal therapy using selective estrogen receptor modulators (SERMs), such as tamoxifen, and aromatase inhibitors. We next investigated whether the candidate targets showing GED were enriched in distinct molecular pathways. Enrichment analyses using Gene Ontology (GO)^31^ for each lineage GED revealed 38 unique pathways enriched across lineages (**Figure 2D**). Although different pathways showed different levels of ED, the two GO terms, (i) “DNA-binding transcription activator activity” (GO:0001216) and (ii) “DNA-binding transcription activator activity, RNA polymerase II-specific” (GO:0001228), were the most frequently-enriched across the lineages (15 out of 18 lineages).

To explore the potential clinical actionability of the identified GEDs, we integrated drug-gene interaction database (DGIdb)^32^, and identified 82 druggable factors out of 244 pan-lineage GEDs (**Figure S3A, Table S3**). By analyzing dependencies at each tissue, we identified 951 druggable targets, in total, showing significant (rho < −0.25, FDR < 0.05) lineage-specific GEDs, including 132 targets for hematopoietic and lymphoid tissue, 101 for lung, 81 for soft tissue, 61 for central nervous system, 52 for ovary, 49 for stomach, 47 for autonomic ganglia, and 44 for breast (**Figure S3B**).

The most strongly-associated (rho ≤ −0.78, FDR < 1e-32) tissue-specific GED targets include MYOD1 in soft tissue, ESR1 in breast, WT1 in ovary, and SOX10 in skin (**Table S3)**. The skin-specific ED observed for SOX10 was also concordant with a recent study^33^, where the mRNA expression of SOX10 was found to be associated with SOX10 hypomethylation and sensitivity to SOX10 knockdown in melanoma cell lines, while other tissue cell lines showed limited SOX10 expression and limited dependency to SOX10 for survival. The strong ESR1-driven dependency in breast cancer cell lines support the established use of SERMs and aromatase inhibitors in ER(+) breast cancers^34^. Several ED genes already have established targeted therapies, and additional gene targets showing strong lineage-specific expression-driven dependencies may also have therapeutic potential.

Based on a set of the most significant GED targets found within lineages (rho < −0.75, FDR < 1e-10), clustering analyses (**Methods**) showed that cancer cells of the pancreas and biliary tract tissue lineages showed the most similar expression-driven dependency profiles, as well as those of the kidney and urinary tract tissue lineages **(Figure S3C**). We also conducted a clustering analysis to identify GED-nominated drug targets showing similar tissue-specificities across tissue lineages. For example, the breast-specific ESR1 transcription factor is clustered with the other factors FOXA1, SPDEF, TBX3, and TRPS1 (**Figure S3D**). These transcription factors showed the strongest GED levels in breast tissue cell lines (rho < −0.6, FDR < 2e-5), where SPDEF showed breast-specific GEDs similar to ESR1. FOXA1 and SPDEF are the key drivers of ER+ breast cancer risk and have been identified as master regulators of the FGFR2-mediated cancer risk^35^. These results identified cross-tissue cancer cells that may share similar targets.

### Cancer vulnerability targets showing protein expression-driven dependency (PED)

Given that gene mRNA expressions show only moderate correlations with protein abundance in cancer^15–20^, we next sought to expand our analyses to identify targets showing protein expression-driven dependency (PED). We applied BEACON to dependency data and protein expression levels in the subset of 290 cell lines with both types of data (**Methods**). BEACON identified 223 proteins showing significant (rho < - 0.25, FDR < 0.05) pan-lineage protein expression-driven dependency (PED). Among the proteins showing pan-lineage PED, just over half (N=123) of the targets also showed significant (rho < −0.25, FDR < 0.05) pan-lineage GED, suggesting general concordance between mRNA and protein while implicating the importance of considering protein expression. ZEB2 was the most strongly-associated PED (rho = - 0.64), followed by FERMT2, GRHL2, KLF5, CDK6, and CCND1 (rho ≤ −0.52), all of which also showed significant GED (**Table S4**, **Figure 3A**, **Figure 3B**). The other 100 PED targets that do not show significant mRNA-level GED included ELMO2, PRDM6, FGFR3, RUNX1, VGLL1, TMEM158, and CBFB (rho ≤ −0.39) (**Table S4**). We identified 78 druggable proteins that show significant (rho < −0.25, FDR < 0.05) pan-lineage expression-driven dependency, including SOX10, MYB, GATA1, MYOD1, CDK6, HNF4A, CCND1, and PAX5 (rho ≤ −0.52) (**Table S5**).

**Figure 3.**
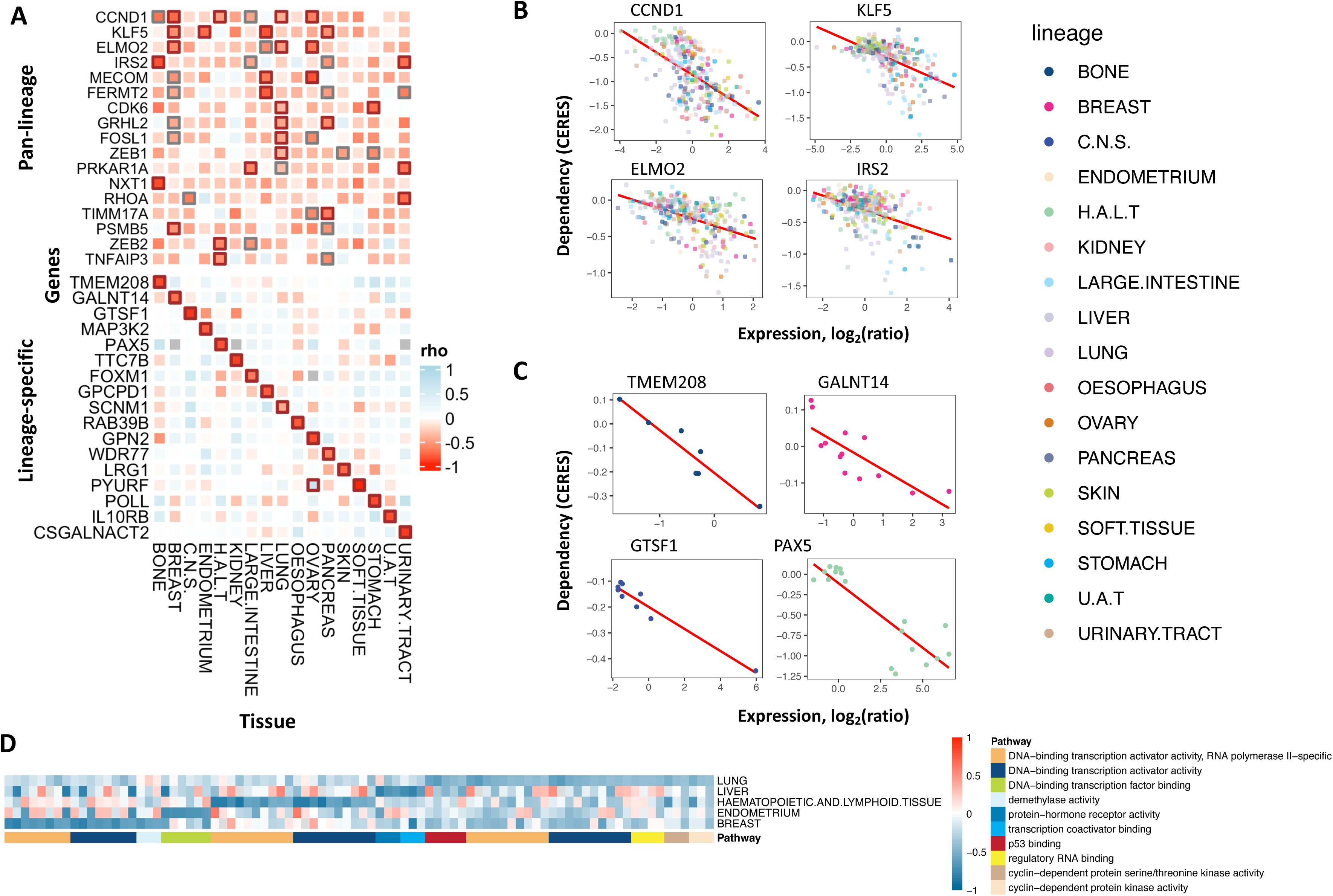
Protein Expression-driven Dependency (PED). (A) Heatmap illustrating pan-lineage and lineage-specific protein expression-driven dependencies (PEDs) across various cancer types. Each square represents the correlation (rho) between protein expression and dependency (CERES scores) in the respective tissue types. Significant dependencies are highlighted with bold outlines (FDR < 0.05 in black, FDR < 0.15 in grey). (B) Scatter plots showing examples of protein expression vs. dependency correlations for selected genes (CCND1, KLF5, ELMO2, IRS2) with significant pan-lineage dependencies. Data points (cell lines) are colored by tissue type. (C) Scatter plots demonstrating lineage-specific dependencies for selected genes (TMEM208, GALNT14, GTSF1, PAX5). Data points are colored by tissue type, highlighting lineage-specific associations. (D) Pathway enrichment analysis of lineage-specific PEDs, visualized as a heatmap. Each cell indicates the ED score of a particular pathway gene (column) in a specific tissue type (row), with genes grouped (colored) by functional pathways.

At the individual tissue level, many of these pan-lineage PED targets also showed high ED within multiple lineages (**Figure 3A**). Targets showing PED exclusive for each lineage (rho ≤ −0.83) included PYURF in soft tissue, GTSF1 in central nervous system, PAX5 in haematopoietic and lymphoid tissue, TTC7B in kidney, and TMEM208 in bone (**Figure 3A**, **Figure 3C, Table S6**). To examine potential actionability of the identified PED proteins, we integrated DGIdb and identified 170 druggable significant (rho < −0.25, FDR < 0.05) lineage-specific PEDs for all lineages; within these, we found a set of very strong lineage-specific targets (rho ≤ −0.81), include PAX5 in haematopoietic and lymphoid tissue, LAMP1 in stomach, NEK6 in urinary tract, TSPO in kidney, CHKA in oesophagus, SERPIND1 in ovary, and SOX10 in central nervous system (**Figure S3E**, **Table S6)**. Enrichment analyses with the PED targets yielded 17 pathways enriched in a more lineage-specific pattern than GED results (**Figure 3D, Table S7**). DNA-binding transcription activator activity (GO:0001216 and GO:0001228) terms were similarly significantly enriched showing consistency with the GED results (3 out of 5 lineages).

### Concordance between gene and protein expression-driven dependency

Protein expression evidence can validate molecular targets observed at the mRNA level. We thus analyze the concordant and unique gene targets based on their GED and PED correlations. We found 123 genes showing consistently significant pan-lineage expression-driven dependency in both mRNA and protein levels, most notably SOX10 (rho_RNA_ = −0.82, rho_protein_ = −0.77), TP63 (rho_RNA_ = −0.69, rho_protein_ = −0.72), IRF4 (rho_RNA_ = −0.82, rho_protein_ = −0.73), and MYB (rho_RNA_ = −0.66, rho_protein_ = −0.75) (**Figure 4A, Table S8**). The confirmation of both GED and PED demonstrate the robustness of these targets.

**Figure 4.**
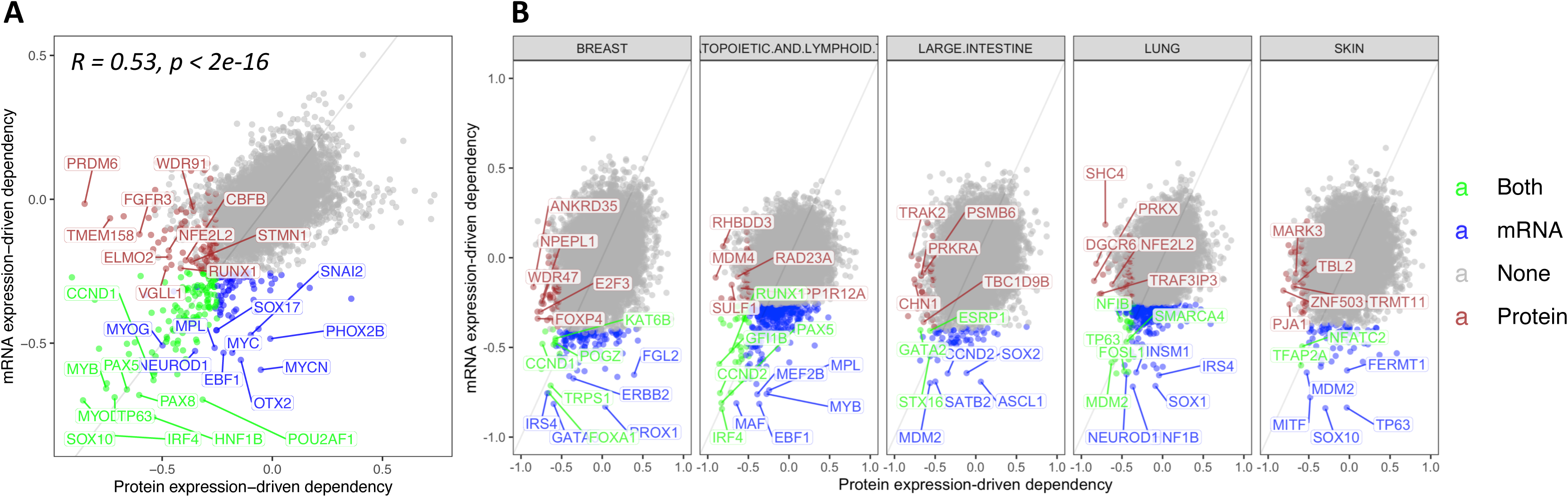
mRNA vs Protein expression-driven dependency. (A) Heatmap illustrating the correlation between pan-lineage GEDs and PEDs across genes. Genes with consistent significant pan-lineage dependencies at both mRNA and protein levels are highlighted, including SOX10, TP63, IRF4, and MYB. Additional significant pan-lineage GEDs without corresponding PEDs (e.g., MYCN, OTX2, EBF1) and PEDs without corresponding GEDs (e.g., ELMO2, PRDM6, FGFR3) are also indicated. (B) Scatter plots showing the correlation between tissue-level GEDs and PEDs within specific lineages.

Meanwhile, given the moderate correlation between mRNA and protein, protein expression-driven dependency may also reveal protein aberrations that arise post-transcriptionally. We found 85 genes showing significant (rho < −0.25, FDR < 0.05) pan-lineage GED without a significant PED that may be less robust as potential therapeutic targets, including MYCN (rho_RNA_ = −0.59, rho_protein_ = −0.05), OTX2 (rho_RNA_ = −0.56, rho_protein_ = −0.14), and EBF1 (rho_RNA_ = −0.53, rho_protein_ = −0.22) (**Figure 4A, Table S8**). On the other hand, we also found 100 proteins showing significant (rho < −0.25, FDR < 0.05) pan-lineage PED without a significant GED. Some notable targets include ELMO2 (rho_RNA_ = −0.2, rho_protein_ = −0.47), PRDM6 (rho_RNA_ = −0.02, rho_protein_ = −0.85), FGFR3 (rho_RNA_ = −0.12, rho_protein_ = −0.6), and RUNX1 (rho_RNA_ = −0.24, rho_protein_ = −0.42) (**Figure 4A, Table S4**).

We next analyzed the consistency between tissue-level GEDs and PEDs for each lineage. (**Figure 4B, Table S9)**. In total, we found 121 genes showing significant GED and PED (rho < −0.25, FDR < 0.05) within a lineage, which may present as some of the strongest targets identified through BEACON. KLF5 showed significant GED and PED (rho < −0.47, FDR < 0.045) in the endometrium, liver, and pancreas lineages. SOX2 gene showed significant GED and PED (rho < −0.42, FDR < 0.032) in the lung and oesophagus lineages. Other lineage-specific targets showing concordance between GED and PEDs include FOXA1 in breast, PAX5 in haematopoietic and lymphoid tissue, GATA2 in large intestine, MDM2/TP63 in lung, and TFAP2A in skin.

### Leveraging expression-driven dependency to enrich for drug targets

Identification of drug targets is a major goal of genomic studies, yet even by using 141,456 human DNA-Seq data in gnomAD without phenotype association, known drug targets only showed a minor difference in constraints for loss-of-function (LoF) variants compared to other genes^28^. To test whether expression-driven dependency derived by BEACON may represent an effective strategy to identify drug targets, we ascertained whether the identified genes showing GED/PED are enriched for druggable targets from DrugBank and curated by Minikel et al.^28^. We used the Fisher’s exact test to evaluate the association between the druggable gene lists and the pan-lineage GEDs/PEDs we identified (**Methods**). The majority (8 out of 15) of druggable gene lists from DrugBank were significantly enriched (Fisher’s exact test, odds ratio > 2, FDR < 0.05) with expression-driven dependency observed in both mRNA and protein level expressions (**Figure 5A, Table S10**). Genes targeted by *Antibody* was the gene set most enriched with GEDs and PEDs (OR_RNA_ = 9.2, OR_protein_ = 18.9), where the higher enrichment in PED align with the mechanism of action of antibody directly binding to proteins. These GED/PED genes include Antibody targets (5 out of 23) showing significant levels of both GED/PED (rho < −0.25, FDR < 0.05) such as CD19 (rho_RNA_ = −0.56, rho_protein_ = −0.66), EGFR (rho_RNA_ = −0.43, rho_protein_ = −0.36), ITGB3 (rho_RNA_ = −0.41, rho_protein_ = −0.36), ERBB2 (rho_RNA_ = −0.41, rho_protein_ = −0.34), and PDGFRA (rho_RNA_ = −0.37, rho_protein_ = - 0.26) (**Figure 5B**, **Figure 5C, Table S11**). These targets also belong to DrugBank’s *Approved drug targets* (OR_RNA_ = 3.9, OR_protein_ = 3.9) and *Oncology (Cancer)* (OR_RNA_ = 10.1, OR_protein_ = 7.5) gene lists, both of which were also significantly enriched with GEDs and PEDs. The high fold enrichment for druggable genes in the *Oncology* gene set align with our analyses using DepMap cancer cell lines. Well-established targets within *Approved drug targets* and *Oncology* that show strong GED and PED include BCL2 (for both RNA and protein level EDs, rho < −0.39), PIK3CD (rho < −0.33), and PDGFRB (rho < −0.29), suggesting. Additionally among the Drugbank *Oncology* gene set, PSMB5 (targeted by proteasome inhibitors bortezomib and carfilzomib for hematologic malignancies) and RXRA (targeted by bexarotene, an RXR agonist used in the treatment of cutaneous T-cell lymphoma (CTCL)), showing significant levels of GED/PED, were among the Oncology gene list, reinforcing the robustness of our approach in identifying clinically relevant targets.

**Figure 5.**
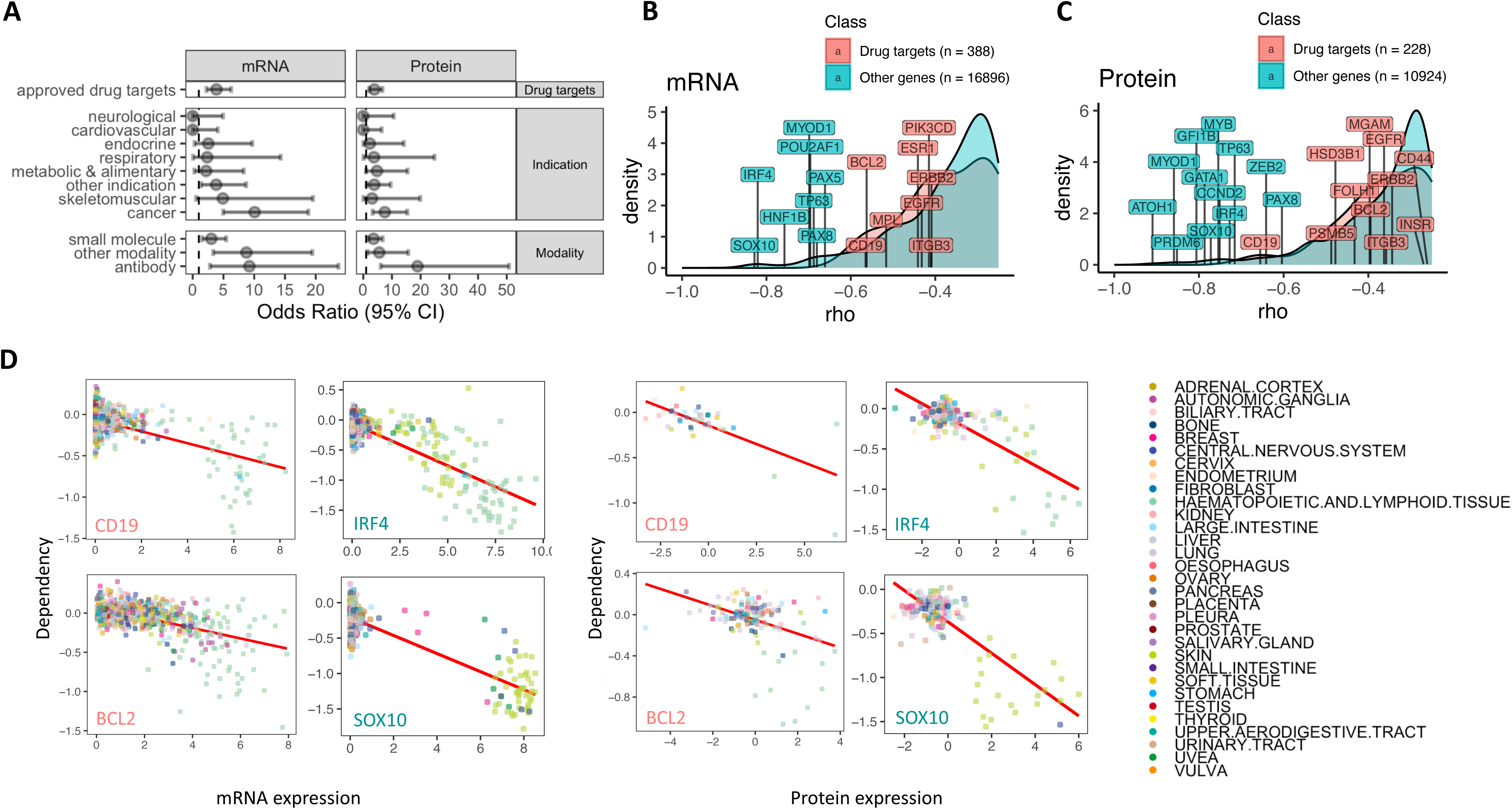
Leveraging Expression-Driven Dependency to Enrich for Drug Targets. (A) Enrichment (Fisher’s exact test) results demonstrating the enrichment of identified GEDs and PEDs in druggable gene lists curated by DrugBank, including all approved drug targets, drug targets by indication, and by drug modality. (B-C) The density plots of ED scores from drug targets (DrugBank approved targets) versus other genes, highlighting the top significant targets identified at (B) mRNA and (C) protein levels. (D) Scatter plots of expression vs. dependency correlations for top drug targets and other genes, showing significant pan-lineage ED at both mRNA and protein levels (e.g., SOX10, TP63, IRF4, CCND1). Data points (cell lines) are colored by tissue type.

Notably, in addition to enrichment for known Oncology druggable genes, BEACON-identified GED/PEDs also showed suggestive enrichment for multiple other indication categories, including DrugBank gene sets for Skeletomuscular (OR = 4.9, p = 0.0693 for GEDs; OR = 3.1, p = 0.29 for PEDs) and Metabolic/Alimentary (OR = 2.2, p = 0.24 for GEDs; OR = 4.80, p = 0.030 for PEDs) diseases. The Drugbank Other Indications category with more targets and statistical power showed significant enrichment for both GEDs (OR = 3.8, FDR =0.022) and PEDs (OR = 3.9, FDR =0.022) for PEDs, suggesting there may be a broader utility of these cell-specific targets beyond oncology.

We next characterized whether GED/PEDs identified by BEACON may be more sensitive to identifying genes with specific mode of inheritance or with additional genetic effect properties^28^. GED/PED genes were both enriched for Autosomal Dominant genes and haploinsufficient genes as determined by ClinGen, but showed no association with Autosomal Recessive genes (**Figure S4**), suggesting these candidates may capture disease genes that are more sensitive to dosage effects. Moreover, GED/PED targets were depleted of genes that were Essential In Culture, confirming that BEACON identifies cell-specific vulnerabilities rather than house-keeping genes that could have off-target effects. Overall, we identified 36 genes in 10 Drugbank/genetic effect lists that showed significant (rho < −0.25, FDR < 0.05) pan-lineage expression-driven dependency in both mRNA and protein levels (**Table S12**).

Additional GED/PED targets identified by BEACON that are not currently druggable targets (DrugBank) include SOX10 (rho_RNA_ = −0.83, rho_protein_ = −0.77, also belong to *ClinGen Haploinsufficient* and *Autosomal Dominant* gene sets) and TP63 (rho_RNA_ = - 0.69, rho_protein_ = −0.72, *ClinGen Haploinsufficient*) and the *Autosomal Dominant* genes GRHL2 (rho_RNA_ = −0.6, rho_protein_ = −0.53) and HNF4A (rho_RNA_ = −0.5, rho_protein_ = −0.62) (**Figure 5B**, **Figure 5C, Table S11**). For genes not in these DrugBank/gene-effect gene lists^28^, BEACON identified 87 targets that showed significant ED (rho < −0.25, FDR < 0.05) at both mRNA and protein levels that may represent potential therapeutic targets for further experimental and clinical development, including IRF4 (for both RNA and protein levels, rho < −0.73), MYB (rho < −0.66), GATA1 (rho < −0.52), FERMT2 (rho < - 0.53), KLF5 (rho < −0.54), CCND1 (rho < −0.54), MYOD1 (rho < −0.7), PAX5 (rho < - 0.66), and CCND2 (rho < −0.64) (**Table S13**). For example, IRF4 knockdown is lethal to multiple myeloma cells^36^. The IRF4 gene is linked to BET protein-mediated transcriptional program^37^ and its dysregulation is also implicated in lymphoid malignancies during hematopoietic cell differentiation^38^. These suggest a therapeutic hypothesis where IRF4-expressing melanoma/lymphoid malignant cells may be accessible through BET inhibitors (BETi).

### Experimental validation of candidate targets showing express-driven dependency

To experimentally validate GED/PED targets identified by BEACON, we selected two types of targets to be tested across two lineages: (1) two targets showing pan-lineage expression-driven dependency, GRHL2 and TP63, and (2) one target showing lineage-specific expression-driven dependency, PAX5. We first confirmed that *TP63* and *GRHL2* mRNA expression were up-regulated in lung squamous (LSCC) tumor tissue compared to tumor-adjacent normal tissue in TCGA, and chose cultured LSCC cells to conduct validation experiments (**Methods**)(**Figure S5A**). After confirming inhibition of target gene expression by shRNA using HARA cells with high expression of the candidate genes (**Figure S5B**), cell proliferation and colony-forming ability were measured using two types of cells with high dependency (HARA, KNS62) on candidate genes and two types of cells with low dependency (H1703, HCC15). In KNS-62 and H1703 LSCC cells, the knockdown of *TP63* using two shRNA constructs (sh-TP63-1 and sh-TP63-2) resulted in a significant reduction in colony formation compared to controls (p<0.01) (**Figure 6A, Figure S5C**). Similarly, *GRHL2* knockdown using sh-GRHL2-1 and sh-GRHL2-2 in both cell lines led to a significant decrease in colony formation (p<0.01) (**Figure 6B, Figure S5D**). *TP63* knockdown also resulted in reduced colony formation in HARA cell line (**Figure S5E**). The results showed that the knockdown of either gene highly inhibited cell viability and colony formation in LSCC cell lines, regardless of the predicted dependence. These results show that pan-lineage targets may represent universal vulnerability and their inhibition may lead to undesired off-target effects on other cells.

**Figure 6.**
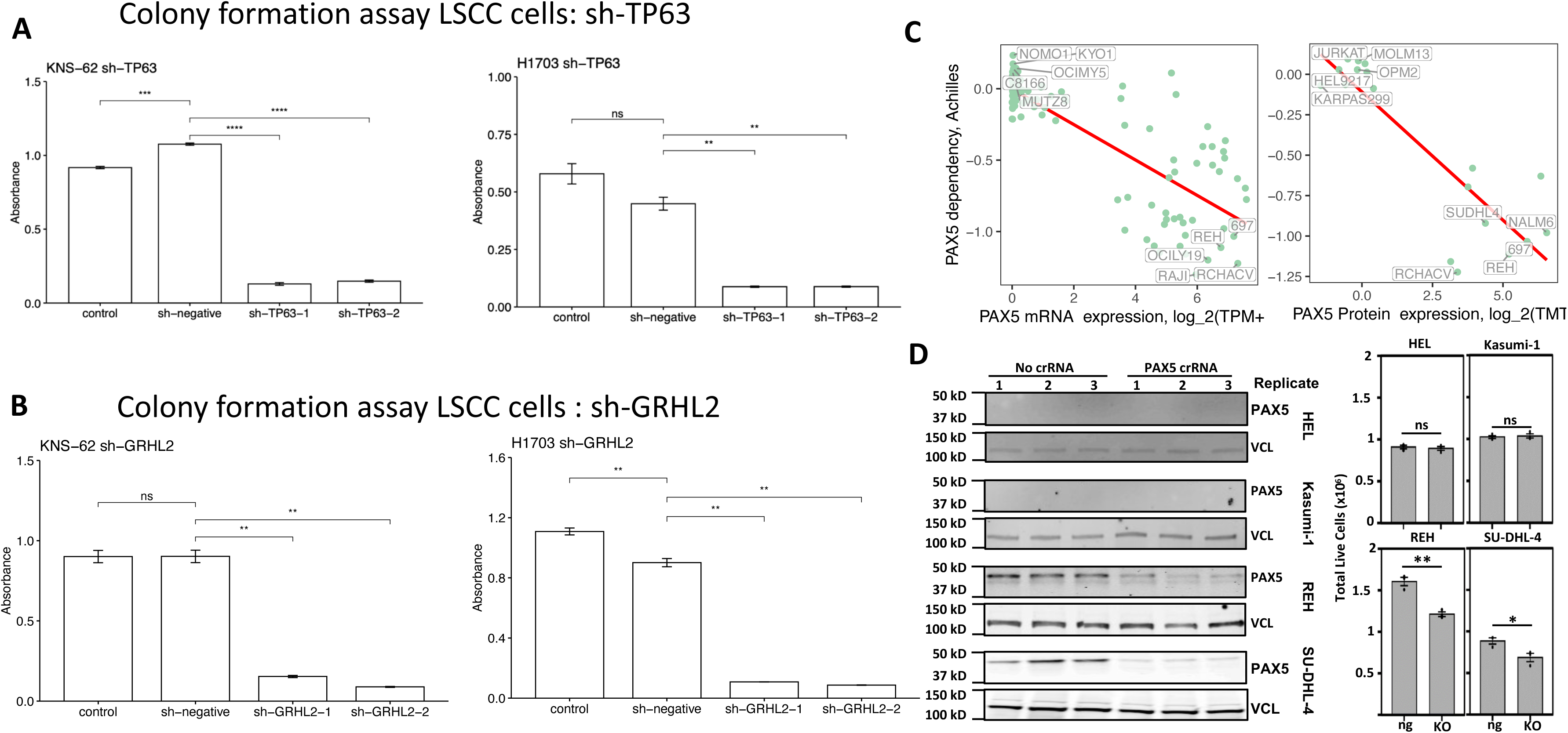
Functional validation of expression-driven dependency targets, TP63, GRHL2, and PAX5, in lung squamous cancer cell and hematopoietic cell lines. (A) Colony formation assay in LSCC cell lines (KNS-62 and H1703) upon knockdown of TP63 using two shRNA constructs (sh-TP63-1 and sh-TP63-2). Significant reduction in colony formation was observed compared to sh-negative control cells (p<0.01). ns: non-significance between control and sh-negative cells. (B) Colony formation assay in LSCC cell lines (KNS-62 and H1703) upon knockdown of GRHL2 using two shRNA constructs (sh-GRHL2-1 and sh-GRHL2-2). Significant decrease in colony formation was seen compared to sh-negative control cells (p<0.01). (C) PAX5 mRNA and protein expression levels in myeloid (HEL, Kasumi-1) and B-cell (REH, SU-DHL4) lineage cell lines, including. PAX5 showed lineage-specific expression-driven dependency. (D) Effect of PAX5 knockout (KO) via CRISPR on cell viability in PAX5-high B-cell lines (REH, SU-DHL4) and PAX5-low myeloid lines (HEL, Kasumi-1). PAX5 KO significantly reduced live cell numbers in REH and SU-DHL4 (p<0.05 and p<0.01, respectively), but not in HEL and Kasumi-1. In (D) left, protein levels were assessed by anti-PAX5 72 hours after electroporation. VCL serves as a loading control. In (D) right, cells were electroporated with RNP complexes with (KO) or without (ng) PAX5 crRNA and allowed to recover for 72 hours. After recovery ng and KO cells were reseeded at equal densities and live cells were counted by trypan blue exclusion after 72 hours.

The lineage-specific target, PAX5, was evaluated for its role in haematopoietic and lymphoid tissue. Within the lineage, groups of cells with high and low PAX5 expression and low and high PAX5 genetic dependency can be clearly identified by BEACON (**Figure 6C**). We chose two PAX5-low myeloid lineage cell lines (HEL and Kasumi-1) and two PAX5-high (REH and SU-DHL4) B-cell lines to conduct PAX5 knockout (KO) experiments via CRISPR. Upon confirming successful KO via western blots, we showed that PAX5 KO significantly reduced the number of live cells in REH and SU-DHL-4 cell lines compared to controls (p<0.05 and p<0.01, respectively). But PAX5 KO did not significantly inhibit cell survival for HEL and Kasumi-1 (**Figure 6D**). Overall, these results show that while TP63 and GRHL2 are essential for cell growth across LSCC cells, PAX5 is specifically crucial for the growth of PAX5-high B cell lymphoma cells. Thus, proteins showing lineage-specific dependencies may present as suitable precision oncology targets in the subset of tumors overexpressing the target gene and protein.

## Discussion

This study integrates large-scale CRISPR screen in conjunction with transcriptomic and proteomic data to identify expression-driven dependencies in cancer cells^4,5^, providing a potential new category of targets in precision oncology, particularly against cancer cells without druggable mutations (**Figure 1**). Our newly developed Bayesian correlation approach BEACON identified known drug targets and uncovered new candidate genes, demonstrating the utility of expression-driven dependency as a complementary strategy to traditional mutation-driven analyses. Functional experiments demonstrated that targeting genes with high expression levels could reveal potential vulnerabilities within specific cancer types, e.g., PAX5 in lymphoid tumors. We also identified distinct molecular pathways enriched in tissues based on the GED/PEDs, providing insights into the biological processes underpinning cancer progression. The concept of expression-driven dependency expands the scope of actionable targets by focusing on genes whose high expression levels selectively contribute to cancer cell survival^7–10^. This is particularly relevant in cases where actionable mutations are absent, thereby addressing a significant gap of treatment options in precision oncology^1–3^.

By integrating CRISPR/transcriptomic data from CCLE/DepMap^4,5^, and global proteomic analyses^24^, we ensured a robust identification of GED/PEDs. GEDs and PEDs show significant correlation (R = 0.54, p < 2e-16) across the cell lines; thus analyzing the GED/PED can cross-validate the robustness of candidate vulnerability targets (**Figure 4**). Our Bayesian approach BEACON further enhanced the reliability of our findings by accommodating variability and limited sample sizes within each tissue lineage (**Figure S2**). The identification of GED/PEDs has significant implications for drug development and personalized cancer therapy. By targeting genes with high expression levels, new therapeutic avenues can be explored in tumors currently with limited treatment options^1–3^. Notably, the strong enrichment of our identified targets with known druggable gene sets highlights the translational potential of our findings (**Figure 5**). While using large human genomic cohort without phenotypes fail to enrich for drug targets^28^, recent human cohort studies demonstrate that genetic evidence provided by genome-wide or mendelian genetic associations can successfully provide 2 to 5 fold enrichment for drug targets^39,40^. We note that our approach here, based solely on data from cell line CRISPR screens, provide an orthogonal approach to refine the drug target search space by providing 3.8 fold enrichment for all drug targets and 7-10 fold enrichment for oncology targets.

We complemented our computational findings with experimental validation. Knockdown of *TP63* and *GRHL2* genes in lung squamous tumor cell lines demonstrated reduced colony-forming ability, and the *PAX5* knock-out cell lines from haematopoietic and lymphoid tissue samples showed reduced cell growth, reinforcing the functional relevance of the vulnerability targets (**Figure 6**). Many GED/PED gene targets are lineage-specific transcription factors; these agree with recent single-cell studies and synthesis that posited the “developmental constraint model of cancer cell states”, which cancer cell states correspond to and may be constrained by the landscape of “developmental map”^41^. Thus, a cancer cell adopting a specific developmental state may require activation of such transcription factors and become genetically dependent. While such targets used to be considered undruggable, new drug modalities such as proteolysis-targeting chimera (PROTAC) are showing strong promises^42–45^.

While our study presents a novel approach to identifying cancer dependencies, several limitations warrant discussion. The reliance on cell line models, despite their widespread use, may not fully capture the complexity of tumor heterogeneity and the tumor microenvironment *in vivo*. Future studies should aim to validate these findings in patient-derived xenografts and clinical samples to confirm their translational potential. Moreover, our Bayesian approach BEACON, while robust (**Figure S2**), is constrained by the quality and completeness of available data (**Figures S1**). Expanding proteomic and transcriptomic datasets that capture the full array of cancer cell heterogeneity across tissue lineages will further improve the reliability of GED/PED identification. Additionally, exploring combination therapies targeting both mutation-driven and expression-driven dependencies could yield synergistic effects, which could be explored in the future.

Overall, our study highlights the potential of expression-driven dependencies as a valuable method for identifying novel therapeutic targets in precision oncology. By integrating multi-omics and CRISPR screen data, we have expanded the repertoire of actionable targets beyond mutated genes for further clinical development, offering new possibilities for cancer treatment.

## Methods

### Data Sources

We used the CCLE mRNA expressions data^33^ and CCLE quantitative proteomics data^24^, and from each dataset we excluded the 26 lineages containing data shared in fewer than 7 cell lines, i.e., Adrenal cortex, Autonomic ganglia, Biliary tract, Brain, Cervix, Colon, Eye, Fibroblast, Melanoma Eye(Skin), Osteosarcoma, Placenta, Pleura, Primary, Prostate, Salivary gland, Skin CJ1(2,3) resistant, Skin FV1(2,3) resistant, Small intestine, Testis, Thyroid, and Uvea. We used the DepMap Public 22Q2 data release from the Cancer Dependency Map Project (DepMap)^5^, which contained the CRISPR knockout screens (Achilles project^25–27^) for 19,221 genes in 1840 cell lines, including both normal and cancer cell lines, corresponding to 33 primary diseases and 30 lineages. We used the druggable gene lists curated in Minikel et al.^28^ The CRISPR knockout screens and mRNA expressions datasets were downloaded from depmap portal^46,47^. The proteomics datasets were downloaded from Nusinow et al.^24^. The druggable gene lists were downloaded from the corresponding studies given in Minikel et al.,^28^ and from the DrugBank resource (release 5.1.7).

### mRNA expression-driven dependency (GED)

To measure the expression-driven dependency of targets we reviewed correlation-based methods utilizing the two variables^48–50^, which are adopted to develop a Bayesian approach that we named BEACON. For each gene, BEACON calculated the Bayesian correlation between the gene’s expressions and CERES cancer dependency scores^25^ across the pan-lineage cell lines. BEACON modeled expression levels and dependency scores as the bivariate Gaussians and used Markov Chain Monte Carlo (MCMC) sampling to estimate the correlation coefficient *rho* between them. Given the null hypothesis that the uncorrelated expression and dependency of a gene has the 0 *rho* coefficient, we statistically tested each gene’s *rho* estimate obtained from the MCMC simulation as follows. Assume that the MCMC sampling is carried out for a null gene’s expression and dependency, then we expect that the distribution of the *rho* estimate accumulated over the MCMC iterations will be centered at zero. Based on this rationale, we computed the z-score of *i-*th gene as the deviation of the MCMC estimate of *rho* from the expected (null) value (i.e., zero) in terms of the standard deviation observed in the simulated distribution, i.e., z(i) = rho_MCMC_(i) / SD_MCMC_(i). Since the z-values, by nature, follow a normal distribution with zero-mean and unit-variance, then we computed the p-value for each gene’s *rho* estimate as the probability of observing a value as extreme as the computed z-value for that gene. We multi-testing corrected the resulting p-values using the BH procedure for FDR. Overall, 4445 genes showed significant pan-lineage expression-driven dependency at the FDR of 0.05. We run the MCMC simulations in R (v3.6) by using *rjags* package (v4-10) with *JAGS* library (v4.3.0).

Compared to other methods that quantify the relationship between two variables, the Bayesian correlation (rho) yielded more intuitive results in the cases with small sample size, while other methods often deviated to spurious correlations imposed by outliers in the data. We benchmarked both methods by simulating expression and dependency datasets at various correlation levels (from −1 to 1, with 0.25 intervals) and sample size (number of cell lines, 10, 20, 30, 40, 60, 100), with different fraction (0.1, 0.3, 0.5) of samples being outliers (**Figure S2**). Through rigorous simulations, we observed that the Bayesian method is better than Pearson correlation for estimating moderate true correlation (|rho| < 0.75) in small sample size (∼10 cell lines). Bayesian method is also preferable in noisy data (noise level ≥ 0.3, i.e., 30% or more of the samples are corrupted by noise to become outliers), regardless of sample size or true correlation level. For large samples (≥60), both methods have similar performance in all settings. Pearson method is only better at detecting fewer (≤20 cell lines) and highly-correlated samples |rho| ≥ 0.75, when there is less noise (≤0.1) (**Figure S2**).

For lineage-wise expression-driven dependency analyses, we stratified by lineage the gene expressions and cancer dependency scores across cell lines, and for each gene we calculated the Bayesian correlation (and the corresponding P value and FDR as in the pan-lineage case) between the gene’s expressions and cancer dependency scores over the lineage cell lines. In median, 270 gene expression-driven dependencies were significant (FDR < 0.05) per lineage. To further identify the lineage-specific targets, we defined a target’s specificity to a given lineage by the difference between the target’s ED score computed within that lineage and the target’s average ED score computed in other lineages.

### Protein expression-driven dependency (PED)

We adopted the aforementioned procedures for analyzing the protein expression-driven dependency. For this, we used the MS proteomics data obtained for 375 cell lines and 22 lineages^24^. For the pan-lineage analysis, we found 907 proteins showing significant expression-driven dependency (FDR < 0.05). For the lineage-wise analyses, we found, in median, 70 proteins per lineage showing significant expression-driven dependency (FDR < 0.05).

### Pathway enrichments from GEDs/PEDs

We used *clusterProfiler*^31^ R package (v4.8.3) for functional enrichment analyses of our identified GED and PED sets, and reported the enrichment GO categories at the BH-adjusted p-value cutoff of 0.05.

### Association of GEDs/PEDs with drug targets

We tested the association between the drug targets and the genes (proteins) that showed significant (rho < −0.25, FDR < 0.05) pan-lineage ED by using the Fisher’s exact test of independence (two-sided). More precisely, given a set of druggable genes, a set of GEDs (PEDs), and the list of total quantified targets in transcriptome (proteome), we calculated the probability of obtaining the observed data and its more extreme deviations in the contingency table consisting of (i) the number of drug targets quantified in the transcriptome (proteome), (ii) the number of GEDs (PEDs) quantified in the transcriptome (proteome), (iii) the number of drug targets quantified in the transcriptome (proteome) that also showed significant ED, and (iv) the remaining number of genes (proteins) that were not drug targets nor showed significant ED, under the null hypothesis that the relative proportions are the same – that the fractions of genes that were drug targets are the same whether the genes show significant ED or not. We found that the pan-lineage GEDs (PEDs) were significantly associated (OR > 2, FDR < 0.05) with the 10 (12) of the druggable gene lists in Minikel at al.^28^

## Methods for the Experimental Validation of TP63 and GRHL2

### Cell culture

The human lung squamous cell carcinoma (LSCC) cell lines, HARA and KNS62 (The Japanese Cancer Research Resource Bank; JCRB, Osaka, Japan), NCI-H1703 and HCC-15 (kindly provided by Dr. John D Minna) were used. KNS62 was cultured in E-MEM culture medium (FUJIFILM Wako Pure Chemical Corporation, Osaka, Japan) containing 20% fetal bovine serum (Sigma-Aldrich Japan, Tokyo, Japan) supplemented with 100 U/mL penicillin and streptomycin sulfate (FUJIFILM Wako Pure Chemical Corporation). The others cell lines were cultured in RPMI-1640 culture medium (FUJIFILM Wako Pure Chemical Corporation) containing 10% fetal bovine serum (Sigma-Aldrich Japan) supplemented with 100 U/mL penicillin and streptomycin sulfate (FUJIFILM Wako Pure Chemical Corporation). All cultured cells were incubated at 37 °C in a humidified atmosphere of 5% CO2 and maintained in continuous exponential growth by passaging. All cell lines were obtained from the reliable biobanks with authentication. Mycoplasma test was performed in regular basis from the first culture of the cells to verify the cells to be the same as the cells registered.

### Plasmid DNA constructs

The shRNA-targeted sequences were listed in **Table S14**. For the constructions of plasmids to express shRNA against target genes, double-stranded oligonucleotides were cloned into the pLKO.1-TRC vector (Addgene, #10878). A nonsense scrambled oligonucleotide was used as a negative control. All of the inserted DNA fragments were confirmed by performing DNA sequencing.

### Lentivirus-mediated transient expression of the constructs in LSCCs

HEK293T cells were transfected with the constructed plasmids along with lentiviral packaging plasmids pVSV-G, pMDL/pPRE and pRSV-REV (Addgene) using a calcium phosphate method. The lentiviral-containing media were collected 72 h after the transfection, filtered through a 0.45 µM filter, then aliquoted and stored at −130°C until use. Cultured LSCC cells were infected with packaged lentiviruses to express shRNA constructs; after 48 hours of culture, the cells were treated with 2.5 μg/ml (HCC15) or 5 μg/ml (HARA, KNS-62, H1703) puromycin (Thermo Fisher, # A1113803) and cultured for 24 hours (HARA, KNS-62, H1703) or 48 hours (HCC15), and used for transient experiments.

### RNA extraction and Quantitative PCR analysis

Gene expression levels were examined by quantitative PCR analysis. Briefly, total RNA was isolated from cells using ISOGEN II (Nippon Gene, #311-07361) and purified using RNeasy Mini Kit (Qiagen). Total RNA (500 ng) was reverse transcribed to cDNA using ReverTra Ace™ (TOYOBO, #FSQ-101). Quantitative PCR was performed using primers listed in **Table S14**, Thunderbird SYBR Green Master Mix (TOYOBO, #QPS-201) and StepOne Plus Real-Time PCR System (Thermo Fisher).

### Cell proliferation and cytotoxic assay

Cell viability was analyzed using the Cell Counting Kit-8 (CCK-8) (Dojindo Laboratories, Kumamoto, Japan: CK04). Cells were seeded 5 x 103/100 μL per well in 96-well plates. After 1 h incubation at 37 °C, 10 μL of CCK-8 solution was added to each well and incubated at 37 °C for 2 h. The absorbance was detected at 450 nm using a plate reader (ThermoFisher Maltiskan FC) according to the manufacturer’s instructions. Cell viability was normalized against the sh-negative control after 24 h of transfection and the data expressed as a ratio against control after 96 h of transfection.

### Colony formation assay

Cells were seeded 1,300-5,000 cells (5,000 cells for HARA, 1,300 cells for KNS-62, 1,500 cells for H1703, 2,000 cells for HCC15) per well into 12 well plates (3.8 cm², Corning Japan, Shizuoka, Japan), and cultured for 10 days with the change of culture media every three days. The cells were then washed by PBS twice, fixed and stained in 0.2% crystal violet dissolved in 20% ethanol, and incubated for 10 minutes at room temperature with gentle shaking. After washing by 1 mL of PBS once and by sterilized water three times, the plate was air dried and photographed. To quantify the colony formation, 1 mL of 50% ethanol (pH 4.2, adjusted by hydrochloric acid) was added into each well of 12-well plates, and incubated for 5 minutes at room temperature with slow shaking, then measured the absorption at 592 nm using a ThermoFisher Maltiskan FC (ThermoFisher). Each experiment was performed with 3 replicate wells.

### Statistical analysis

Data were analyzed using R version 4.0.3 (The R Foundation for Statistical Computing, Vienna, Austria) in combination with R studio version 1.2.5033 (R studio, Boston, MA, USA). Welch two sample t-test was used to examine statistical difference between two groups.

## Methods for the Experimental Validation of PAX5

### Tissue culture

All cell lines used in this study were maintained at 37°C with 5% CO^2^. HEL and SU-DHL4 cells were cultured in RPMI 1640 (Corning) supplemented with 10% FetalPlex serum (Gemini) 1% L-Glutamine (Corning), and 1% Penicillin Streptomycin (Gibco). Kasumi-1 cells were cultured in RPMI 1640 (Corning) supplemented with 15% FetalPlex serum (Gemini) 1% L-Glutamine (Corning), and 1% Penicillin Streptomycin (Gibco). REH cells were cultured in IMDM (Gibco) supplemented with 10% heat inactivated FBS (R&D Systems), 1% L-Glutamine (Corning), and 1% Penicillin Streptomycin (Gibco).

### Genome Editing

The CRISPR/Cas9 system was used to genetically engineer cell lines via ribonucleoprotein (RNP) complex delivery as previously described (Layden et al., 2021). Briefly, a crRNA (IDT) targeted to exon 4 of PAX5, CTTTTGTCCGGATGATCCTG, was annealed with tracrRNA (IDT). Control RNP complexes were formed without the crRNA. Annealed gRNA were incubated with S.p. Cas9 Nuclease (IDT) to form RNP complexes and electroporated into 1.25 million cells per condition using the NEON transfection system (ThermoFisher). Cells were grown for 72 hours and knockout efficiency was assessed by western blot. Electroporations were performed in biological triplicate for each condition.

### Growth Analysis

Cells were allowed to recover for 72 hours post electroporation and then were reseeded to 0.2-0.5 x 10^6^ depending on cell line. Reseeded cells were incubated for 72 hours at 37°C with 5% CO2. Cells were mixed with Trypan Blue (Gibco) and counted with a hemocytometer. Cells were counted in technical triplicate for each biological replicate.

### Western Blots

Protein was isolated from cells lysed with RIPA buffer (50 mM Tris pH 8.0, 150 mM NaCl, 1% NP-40, 0.5% sodium deoxycholate, 0.1% SDS) and sonicated before centrifugation. Protein concentration was quantified using the DC Protein Assay kit (BioRad). Equal amounts of protein were boiled in Laemmli buffer and run on SDS-PAGE gels. Proteins were transferred to a PVDF membrane and membranes were blocked in 5% BSA in PBS. Membranes were incubated with the indicated primary antibodies diluted in 5% BSA in PBS-T followed by incubation with IRDye 800CW and 680RD secondary antibodies (LI-COR Biosciences) diluted in PBS-T + 0.01% SDS. Blots were imaged using the Odyssey Imaging System (LI-COR Biosciences). The following primary antibodies were used at a 1:1000 dilution: PAX5 (Santa Cruz; A-11) and VCL (Santa Cruz; 7F9). All secondary antibodies (IRDye 680 Donkey anti-Rabbit and IRDye 800 Donkey anti-Mouse, Licor) were used at a 1:5000 dilution.

### Statistics

Statistical analyses were performed in R (version 4.1.0). Unpaired two-sample t-tests were used to determine significance between conditions.

**Supplementary Data**. Spreadsheets containing the supplementary tables (S1-S14), including the related data used for plotting main figures and supplementary figures.

**Table S1.** Data availability for all lineage cell lines.

**Table S2.** BEACON results of pan-lineage GEDs (tissue-specific GEDs – sheet2).

**Table S3.** BEACON results of pan-lineage druggable GEDs (tissue-specific druggable GEDs – sheet2).

**Table S4.** Proteins showing significant (rho < −0.25, FDR < 0.05) pan-lineage PED with significant GED (without a significant GED – sheet2).

**Table S5.** BEACON results of pan-lineage druggable PEDs.

**Table S6.** BEACON results of tissue-specific PEDs (tissue-specific druggable PEDs – sheet2).

**Table S7.** Gene ontology enrichment analysis for the tissue-specific PEDs.

**Table S8.** Genes showing consistently significant (rho < −0.25, FDR < 0.05) pan-lineage expression-driven dependency in both mRNA and protein levels (in only mRNA levels – sheet2).

**Table S9.** Table showing consistency between tissue-level GEDs and PEDs for each lineage.

**Table S10.** Results of Fisher’s exact test evaluating the association between the druggable gene lists and the pan-lineage GEDs/PEDs.

**Table S11.** GEDs/PEDs enriched for druggable targets curated by the DrugBank.

**Table S12.** Druggable genes significantly enriched with expression-driven dependency observed in both mRNA and protein level expressions.

**Table S13.** Other genes (not drug targets) significantly enriched with expression-driven dependency observed in both mRNA and protein level expressions

**Table S14.** The shRNA-targeted sequences and primers used for LSCC experiments.

## DATA AND SOFTWARE AVAILABILITY

### Data Availability

Data for DepMap/CCLE genetic screens and mRNA/protein expressions can be found on DepMap data portal: https://depmap.org/portal/.

### Code Availability

The source code for BEACON is available at https://github.com/Huang-lab/BEACON.

## ACKNOWLEDGEMENTS

The authors wish to acknowledge data from the Cancer Dependency Map project. The authors thank all members of the Huang lab for constructive discussion. Large language models (LLM) may have been used in the initial drafts of coding and writing of this work. All final codes and texts have been extensively edited and verified by the authors. This work was supported by NIH NIGMS R35GM138113, ACS RSG-22-115-01-DMC to KH.

## CONTRIBUTIONS

KH conceived the research, and AE and KH designed the approach and computational analyses. AE developed the Bayesian approach and conducted the bioinformatics analyses, which is supervised by KH. HL, JE, LB, XZ, RI, HO, and SH designed and conducted the experiments. AE, HL, SH, and KH wrote the manuscript. All authors read, edited, and approved the manuscript.

### Declaration of generative AI and AI-assisted technologies in the writing process

During the preparation of this work the authors used ChatGPT, Perplexity, and Claude in order to refine language and assist with editing the authors originally written content for improved readability. After using this tool/service, the authors reviewed and edited the content as needed and take full responsibility for the content of the publication.

