## Supplementary Figures for "Expression-Driven Genetic Dependency Reveals Targets for Precision Medicine"

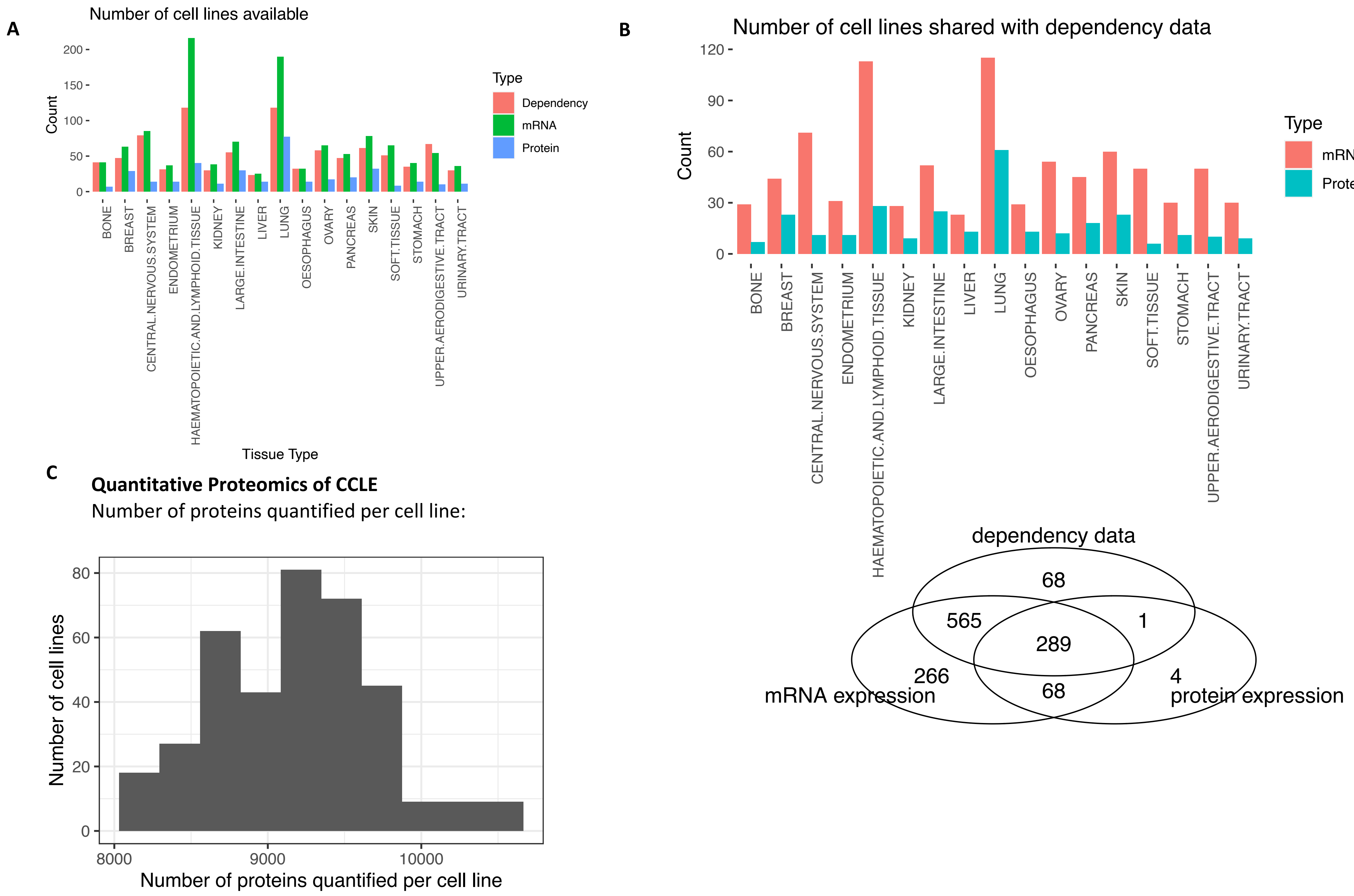


**Figure S1. Data overview.** (A) Analyses were restricted to lineages with at least 7 cell lines having cancer cell line dependency and corresponding mRNA/protein expression data to ensure statistical robustness. (B) 855 cell lines across 17 lineages were analyzed, sharing cancer cell dependency scores and corresponding mRNA and protein expressions. The limited sample size per cell lineage may lead to spurious correlations, especially for protein expression. (C) The distribution of protein quantification per cell line. Over 12,000 proteins (in total) were quantified across all samples, where a majority of the samples reached a quantification level of over 9,000 proteins (PMID: 31978347).

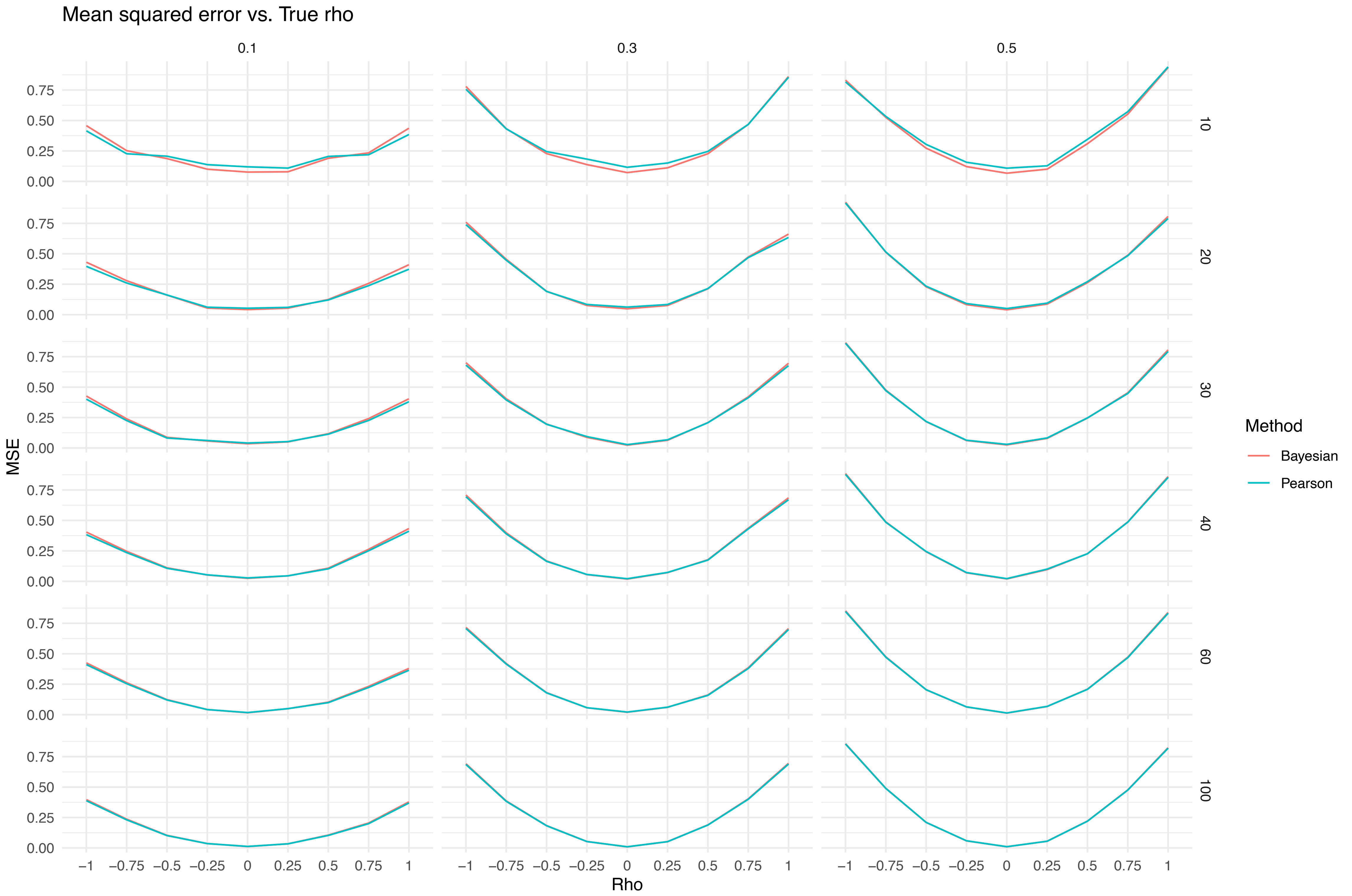


**Figure S2. Benchmarking Bayesian correlation and Pearson's method.** The performance are measured by mean squared-error (MSE, y-axis) for the same data sets randomly simulated for various true correlation levels (rho, x-axis), under different conditions of noise interference (columns, 0.1, 0.3, 0.5) and sample size (rows, 10, 20, 30, 40, 60, 100).

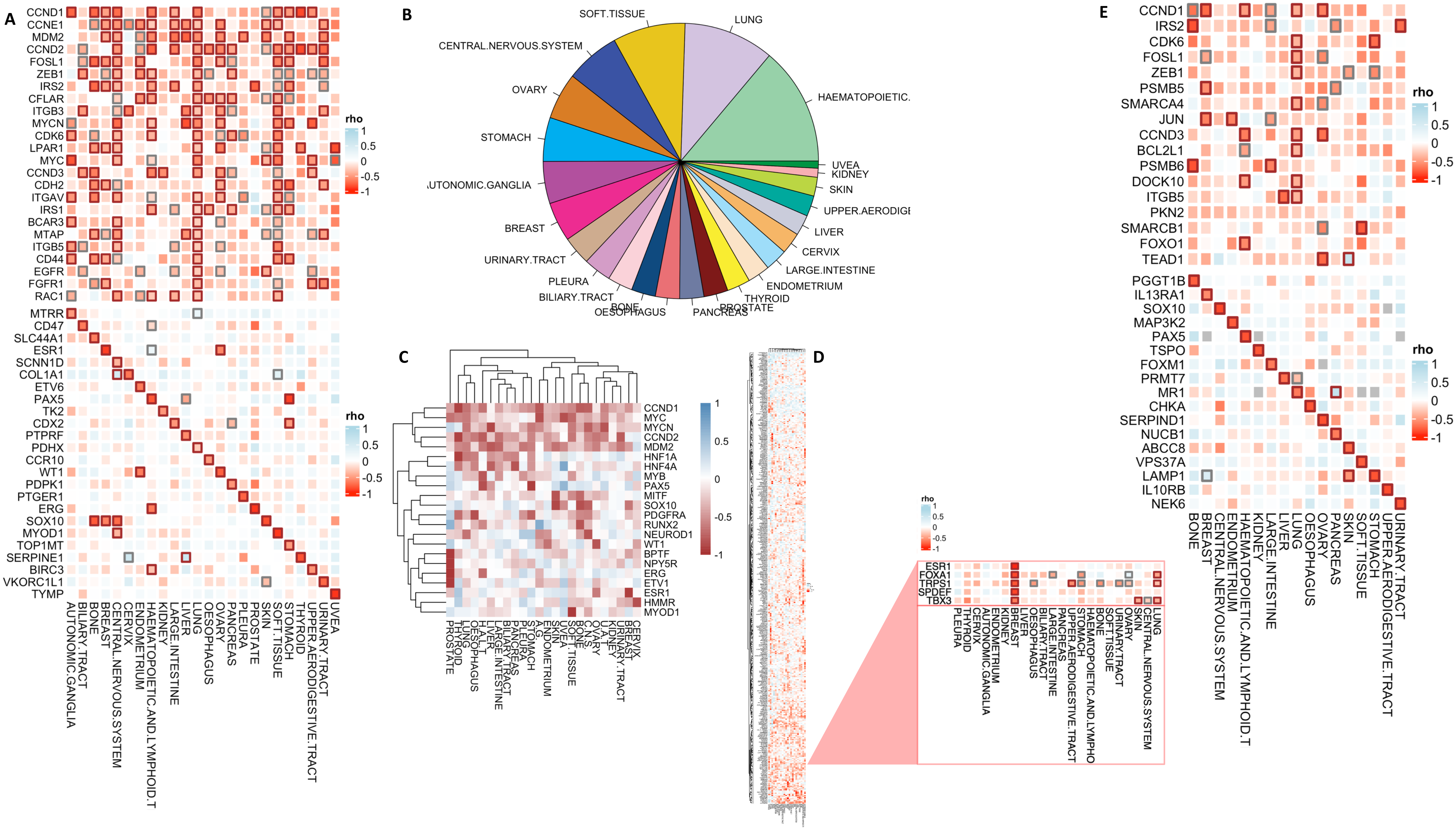

**Figure S3. Analysis of Druggable Gene Expression Dependencies (GEDs) and Protein Expression Dependencies (PEDs).** (A) Heatmap illustrating pan-lineage and lineage-specific druggable gene expression-driven dependencies (GEDs) across various cancer types. Each square represents the correlation (rho) between gene expression and dependency (CERES scores) in the respective tissue types. Significant dependencies are highlighted with bold outlines (FDR < 0.05 in black, FDR < 0.15 in grey). Integration of the drug-gene interaction database (DGIdb) identified 82 druggable factors showing pan-lineage GED and 951 tissue-specific druggable targets, in total, showing significant GED across all lineages. (B) Analysis of tissue-specific expression-driven dependencies across tissues revealed 951 significant druggable targets, including 132 for hematopoietic and lymphoid tissue, and 101 for lung. (C) Clustering GED measures of druggable genes across tissue types showed that pancreatic and biliary tract cancer cells, as well as kidney and urinary tract cancer cells, have similar druggable profiles. (D) The breast-specific ESR1 transcription factor clustered with other factors such as FOXA1, SPDEF, TBX3, and TRPS1, showing strong GED levels in breast tissue cell lines. (E) Integration of DGIdb for PEDs identified 170 significant lineage-specific PEDs, with notable targets including PAX5 in hematopoietic and lymphoid tissue, and SOX10 in the central nervous system.

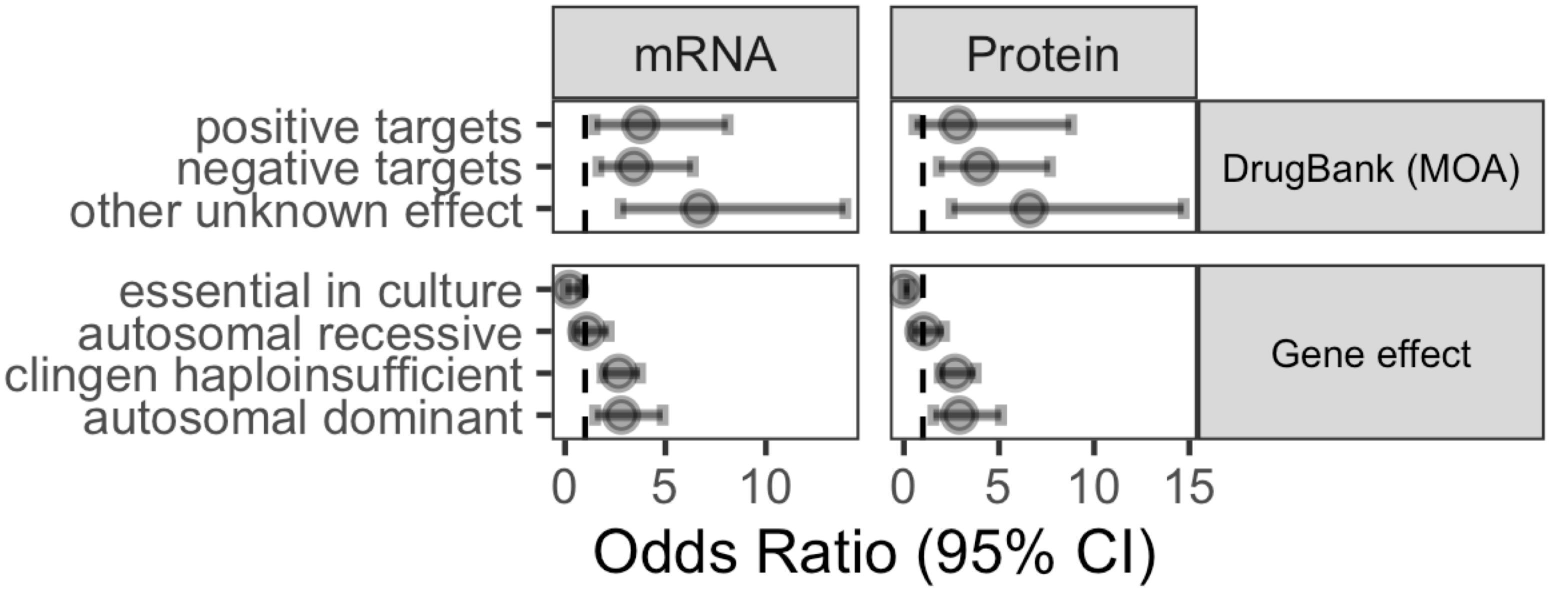


**Figure S4. Enrichment (Fisher's exact test) results demonstrating the enrichment of identified GEDs and PEDs in druggable gene lists based on DrugBank (likely mechanism of action) and genetic effect gene lists as described in Methods.**

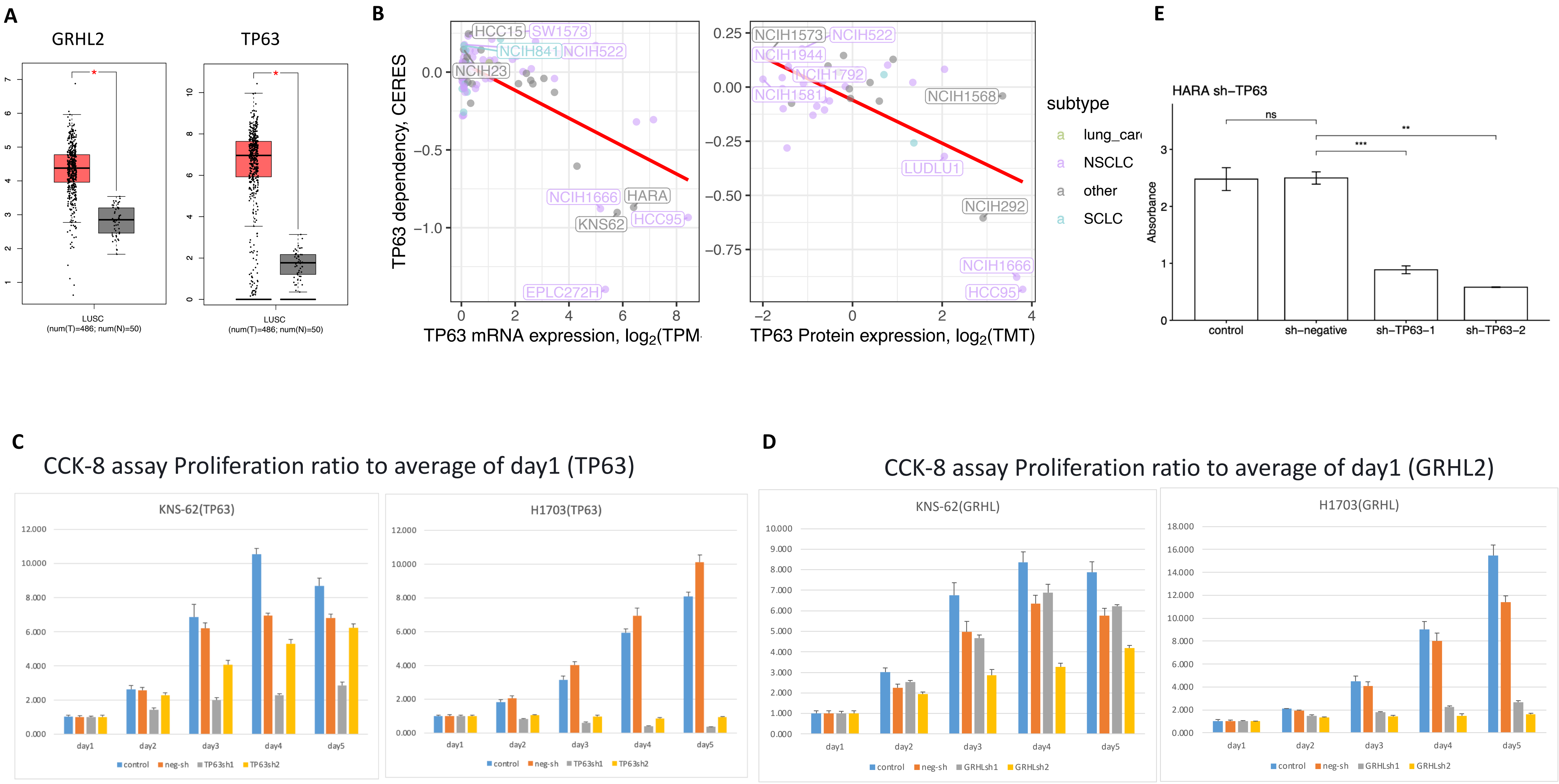


**Figure S5. Dependencies of GRHL2 and TP63 in lung squamous cell carcinoma (LSCC) cell lines.** (A) mRNA expression levels of GRHL2 and TP63 in TCGA LUSC tumor vs. normal tissues in TCGA, where both genes show elevated expression in LSCC compared to normal adjacent normal lung tissue. (B) Scatter plots showing the relationship between TP63 mRNA and protein expression levels vs. gene dependency (CERES score) across various cell lines. (C) CCK-8 cell proliferation assays in LSCC cell lines (KNS-62 and H1703) upon knockdown of TP63 using two shRNA constructs (sh-TP63-1 and sh-TP63-2). Proliferation is shown as a ratio to the average of day 1. (D) CCK-8 cell proliferation assays in LSCC cell lines (KNS-62 and H1703) upon knockdown of GRHL2 using two shRNA constructs (sh-GRHL2-1 and sh-GRHL2-2). Proliferation is shown as a ratio to the average of day 1. (E) Colony formation assay in HARA LSCC cells upon TP63 knockdown. Significant reduction in colony formation is observed compared to control.
